# Prmt5 promotes vascular morphogenesis independently of its methyltransferase activity

**DOI:** 10.1101/2020.02.27.967810

**Authors:** Aurélie Quillien, Manon Boulet, Séverine Ethuin, Laurence Vandel

**Affiliations:** Centre de Biologie du Développement (CBD), Centre de Biologie Intégrative (CBI), Université de Toulouse, CNRS, UPS, France; Université Clermont Auvergne, CNRS, Inserm, GReD, F-3000 Clermont-Ferrand, France

**Author notes:** Corresponding Authors Aurélie Quillien Laurence Vandel.

**Keywords:** Prmt5, zebrafish, angiogenesis, hematopoiesis, endothelial cells, chromatin looping

## Abstract

During development, the vertebrate vasculature undergoes major growth and remodeling. While the transcriptional cascade underlying blood vessel formation starts to be better characterized, little is known concerning the role and mode of action of epigenetic enzymes during this process. Here, we explored the role of the Protein Arginine Methyl Transferase Prmt5 during blood vessel formation and hematopoiesis in zebrafish. Through the generation of a *prmt5* mutant, we highlighted a key role of Prmt5 in both hematopoiesis and blood vessel formation. Notably, we showed that Prmt5 promotes vascular morphogenesis through the transcriptional control of ETS transcription factors and adhesion proteins in endothelial cells. Interestingly, we found that Prmt5 methyltransferase activity is not required to regulate gene expression, and the comparison of chromatin architecture impact on reporter genes expression leads us to propose that Prmt5 rather regulates transcription by acting as a scaffold protein that facilitates chromatin looping in these cells.

## INTRODUCTION

Blood vessel formation is an essential developmental process required for the survival of all vertebrates and much effort has been devoted to understand the molecular pathways and to identify key molecules that regulate different aspects of this process. Interestingly, the vascular anatomy and the mechanisms involved in vessel formation are highly conserved among vertebrates (for a review, (Isogai et al., 2001)). Hence, in the past two decades, zebrafish has been proven to be a useful model to study vascular morphogenesis and blood cell formation *in vivo* (Beis and Stainier, 2006; Lawson and Weinstein, 2002a; Thisse and Zon, 2002).

In vertebrates, blood cell formation is tightly associated with the development of the vascular system. Hematopoietic Stem Cells (HSC), which give rise to the different blood cell lineages, emerge directly from the ventral part of the dorsal aorta, an area referred to as the hemogenic endothelium. Notably, the ETS transcription factor ETV2 functions as a master regulator for the formation of endothelial and hematopoietic cell lineages through the induction of both blood cells and vasculature transcriptional programs, in mouse and in zebrafish (Liu et al., 2015b; Wong et al., 2009). In endothelial cells, ETV2 regulates the expression of other ETS transcription factors, VEGF (Vascular Endothelial Growth factor) signaling receptors and effectors, Rho-GTPases and adhesion molecules (Liu et al., 2015b; Wong et al., 2009). Besides, adhesion molecules have been shown to be crucial players in vascular morphogenesis as Vascular Endothelial cadherin (VE-cad/ cdh5) and endothelial cell-selective adhesion molecule (Esama) are essential for junction remodeling and blood vessel elongation in zebrafish (Sauteur et al., 2017; Sauteur et al., 2014). Indeed, loss of function of both *cdh5* and *esama* leads to the formation of disconnected vessels and delayed lumen formation. Likewise, knock down of the scaffold protein Amolt2, which associates to VE-cadherin, also leads to sprout elongation defects and narrowed aortic lumen (Hultin et al., 2014). While the transcriptional cascade underlying blood vessel formation starts to be better characterized, little is known concerning the role and mode of action of epigenetic enzymes during this process. Even though chromatin-modifying enzymes have been described as central in cardiovascular disease and development (Rosa-Garrido et al., 2018; Shailesh et al., 2018), only few examples illustrate in detail the role of epigenetic enzymes during blood vessel development. For instance, the chromatin-remodeling enzyme BRG1 affects early vascular development as well as hematopoiesis in mice (Griffin et al., 2008), and the histone acetyltransferase P300 has been proposed to be recruited at the promoter of specific endothelial genes by the ETS transcription factor ERG (ETS Related Gene) to control their expression both *in vivo* in zebrafish and in HUVEC (Human Umbilical Vein Endothelial Cell) (Fish et al., 2017; Kalna et al., 2019).

Given the common origin of blood and endothelial cells, and their partially shared transcriptional programs, it is plausible that known chromatin-modifying enzymes affecting hematopoiesis could also control blood vessel formation. Along this line, the epigenetic enzyme Prmt5 (Protein Arginine Methyltransferase 5) has been identified as a key player in blood cell formation (Liu et al., 2015a) but its impact on endothelial development has not been investigated to date. Prmt5 catalyzes the symmetric di-methylation of arginine residues on a variety of proteins including histones and therefore acts on many cellular processes such as genome organization, transcription, differentiation, cell cycle regulation or spliceosome assembly, among others (Blanc and Richard, 2017; Karkhanis et al., 2011; Stopa et al., 2015). Prmt5 is mainly known to repress transcription through the methylation of arginine residues on histones H3 and H4 and has been shown to regulate several differentiation processes such as myogenesis, oligodendrocyte and germ cell differentiation or hematopoiesis (Batut et al., 2011; Liu et al., 2015a; Shailesh et al., 2018; Zhu et al., 2019). In mice, *prmt5* knock out prevents pluripotent cells to form from the inner cell mass and is embryonic lethal (Tee et al., 2010). Conditional loss of *prmt5* in mice leads to severe anemia and pancytopenia and Prmt5 maintains Hematopoietic Stem Cells (HSCs) and ensures proper blood cell progenitor expansion (Liu et al., 2015a). Loss of *prmt5* leads to oxidative DNA damages, increased cell apoptosis due to p53 dysregulation and as a consequence, to HSC exhaustion. In this context, Prmt5 protects HSCs from DNA damages by allowing the splicing of genes involved in DNA repair (Tan et al., 2019).

Here, we explored the role of the Protein Arginine MethylTransferase Prmt5 during blood vessel formation and in hematopoiesis in zebrafish. Through the generation of a *prmt5* mutant, we highlight the key role of this gene during vascular morphogenesis *via* the control of expression of several ETS transcription factors and adhesion molecules. Moreover, we show that Prmt5 methyltransferase activity is not required for blood vessel formation and our results suggest that Prmt5 helps to shape correct chromatin conformation in endothelial cells.

## RESULTS

### Prmt5 is required for HSC maintenance and lymphoid progenitor expansion

To characterize *prmt5* function, we generated a *prmt5* mutant by targeting the second exon of *prmt5* with the CRISPR/Cas9 system. A deletion of 23 nucleotides was obtained, leading to a premature stop codon before the catalytic domain of Prmt5 (Fig. 1A). As a consequence, Prmt5, which was expressed ubiquitously in the trunk at 24 hours post fertilization (hpf), was no longer detected in the mutant (Fig. 1B, C). Similarly, Prmt5 expression was severely reduced in *prmt5* morpholino-injected embryos as compared to control morphants (Fig. S1 A, B) (Batut et al., 2011). In order to test whether Prmt5 regulates hematopoiesis in zebrafish, we took advantage of the transgenic line *Tg*(*gata2b:Gal4;UAS:lifeactGFP)* that labels Hematopoietic Stem Cells (HSCs) (Butko et al., 2015). HSCs emerge from the ventral wall of the dorsal aorta (DA, Fig. 1D, D’), before migrating into the Caudal Hematopoietic Tissue (CHT) (Fig. 1D) where Hematopoietic Stem and Progenitor Cells (HSPCs) proliferate and undergo maturation (Butko et al., 2015). Reminiscent of the data published in mice (Liu et al., 2015a), the loss of *prmt5* led to an increased number of *gata2b+* HSCs in 36 hpf mutant embryos as compared to wild type ones (Fig. 1E-G). In addition, we found that the relative expression of *scla*, *runx1* or *cmyb,* which are specifically expressed in emerging HSCs, was increased in *prmt5* mutant embryos as compared to wild type embryos (Fig. 1H). These results suggest that Prmt5 regulates the number of emerging HSCs from the dorsal aorta. We next investigated whether blood cell formation was impaired in *prmt5* zebrafish mutant as described in mouse (Liu et al., 2015a). HSPCs give rise to different blood cell progenitors, such as lymphoid progenitors which colonize the thymus leading to T lymphopoiesis (Fig. 1D) (Ma et al., 2013). As *gata2b+* lymphoid progenitors deriving from *gata2b+* HSCs can be detected in the thymus of transgenic zebrafish larvae from day 3 (Butko et al., 2015), we investigated whether Prmt5 could act on theses progenitors. Indeed, we found that at 5 days, the number of *gata2b+ lymphoid* progenitors in the thymus was significantly reduced in *prmt5* mutant and in morphant embryos as compared to wild type embryos (Fig. 1I-K, Fig. S1 C, D, G), suggesting that Prmt5 is required for lymphoid progenitor expansion. Altogether, these data indicate an important and conserved role of Prmt5 during hematopoiesis in zebrafish as in mouse.

**Figure 1:**
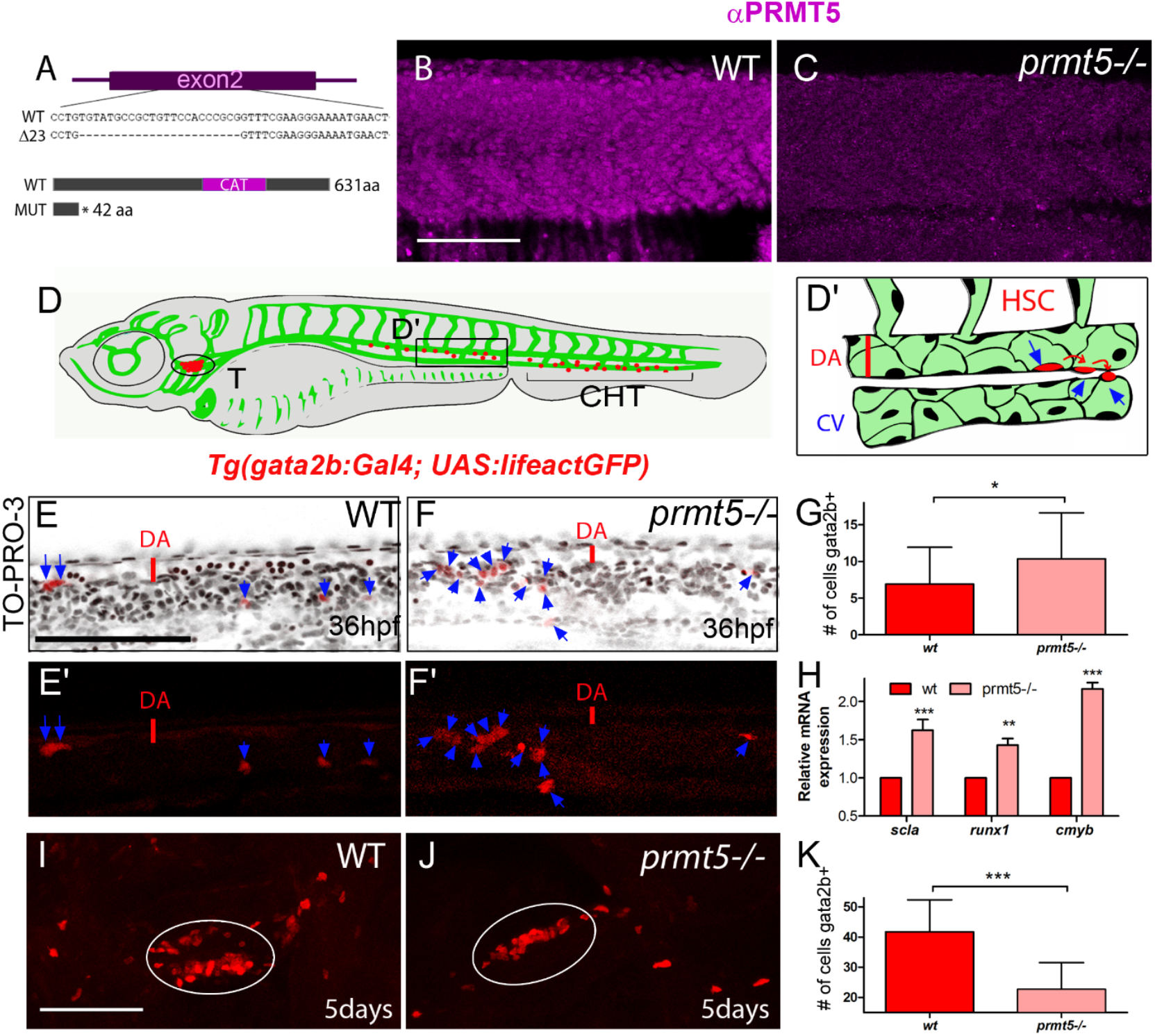
Loss of *prmt5* affect HSCs and HSPCs production. **A**-Schematic representation of the sequence targeted by CRISPR/Cas9 leading to a 23 nucleotides deletion, and of wild type and truncated Prmt5 proteins. The catalytic domain “CAT” appears in magenta. **B**-**C**-Confocal sections of immunostaining with anti-Prmt5 antibody of wild type and *prmt5* mutant embryos at 24 hpf. Scale bar 100 μm. **D**-Schematic representation of vascular (green) and hematopoietic (red) systems in a zebrafish larva. Circle and bracket indicate the Thymus (T) and the Caudal Hematopoietic Tissue (CHT), respectively. **D’**-Close-up of the trunk vasculature where HSCs emerge from the ventral wall of the dorsal aorta (DA), bud and migrate. Red line represents the diameter of the dorsal aorta. Cardinal Vein (CV). **E**-**F’**-Confocal section of transgenic *Tg*(*gata2b:Gal4; UAS:lifeactGFP*) embryos at 36 hpf showing gata2b+ cells in red and TO-PRO-3 in black. Blue arrows indicate HSCs labelled in red in wild type (**E**, **E’**) and in *prmt5* mutant (**F**, **F’**) embryos. Bar scale 100 μm. **G**-Average number of HSCs enumerated per confocal stack in wild type and in *prmt5* mutant embryos at 36 hpf. Data are from 3 independent experiments with at least 6 individuals per experiment and a Mann-Whitney test was performed. **H**-Relative mRNA expressions determined by RT-qPCR in 36 hpf wild type and *prmt5* mutant embryos, from 3 independent experiments with at least 6 animals per condition. Two-way ANOVA was performed. **I**-**J**-Confocal sections of wild type (**I**) and *prmt5* mutant (**J**) thymus from transgenic Tg(*gata2b:Gal4; UAS:lifeactGFP*) embryos at 5 days. Thymus are delimited by a white circle. Bar scale 100 μm. **K**-Average number of HSPCs enumerated per confocal stack in wild type and *prmt5* mutant embryos at 5 days from 3 independent experiments with at least 5 individuals per analysis. T-test was performed. * P<0.05, ** P<0.01, ***P<0.001.

### Prmt5 is required for vascular morphogenesis

As Prmt5 regulates zebrafish hematopoiesis, we next asked whether Prmt5 could also play a role during blood vessel formation, either during angiogenesis or vasculogenesis. First, we analyzed the expression and localization of Prmt5 by immunostaining in *Tg(fli1a:eGFP)* transgenic embryos, in which endothelial cells can be visualized with *egfp* (Lawson and Weinstein, 2002b). We found that Prmt5 was clearly expressed in early endothelial cells at 14 somite stage (Fig. 2A-A’’). At 24 hpf, Prmt5 was expressed in endothelial cells of the dorsal aorta (DA) and of the cardinal vein (CV) (Fig. 2B, B’, D). Prmt5 was also detected in Intersegmental Vessels (ISVs) sprouting from the DA, in either the tip cell (leading the sprout) or the stalk cell (Fig. 2C, C’, D). We then analyzed whether blood vessel formation was affected in transgenic *Tg(fli1a:eGFP) prmt5* mutants at 28 hpf. We found that the dorsal aorta diameter of mutant embryos was reduced as compared to the control (Fig. 2D, E, F close-ups), suggesting that lumen formation was perturbed. To confirm this result, we made use of the Notch reporter line *Tg(TP1bglob:VenusPEST)^s940^* in which only the dorsal aorta cells express the transgene while the cardinal vein endothelial cells do not (Ninov et al., 2012; Quillien et al., 2014). In this transgenic context the area occupied by the dorsal aorta in *prmt5* morphant embryos was significantly reduced as compared to control embryos (Fig. 2G-I). *Prmt5* mutant embryos also showed a defect of sprouting ISV to reach the most dorsal part of the trunk and to connect with other ISVs and form the Dorsal Longitudinal Anastomotic Vessel (DLAV) (Fig. 2D, E, F). This defect was associated with a significant reduction of ISV length (Fig. 2E, F, K) but with no impact on the number of endothelial cells (Fig. 2J). The observed size reduction of ISVs is thus most likely the result of an elongation issue rather than a proliferation defect. Of note, *prmt5* morphants reproduced the phenotype observed in *prmt5* mutants *i.e.* a reduced ISV length at 28 hpf (Fig. S2 A-D).

**Figure 2:**
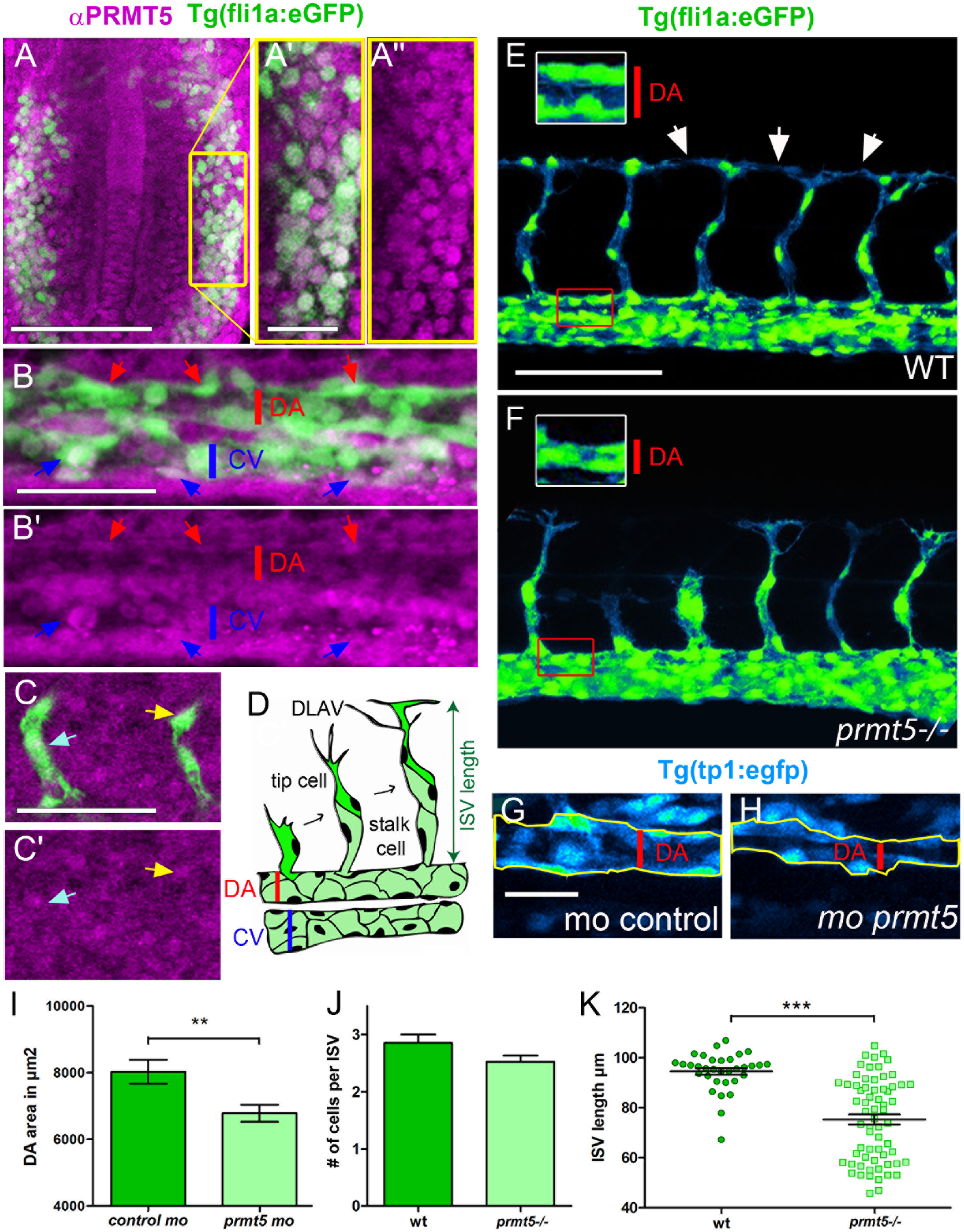
Loss of *prmt5* impairs blood vessel formation. **A**-**C’**-Confocal projections of transgenic *Tg(fli1a:GFP)^y1^* embryos with endothelial cells (in green) after immunostaining against Prmt5 (in magenta). **A**-**A’’**-Dorsal view of the lateral plate mesoderm at 14 somite-stage. Yellow rectangle delimits the close up of Prmt5+ endothelial cells (**A’**-**A’’**). Prmt5+ cells appear in magenta (**A**-**A”**) and endothelial cells in green (**A**-**A’**). Anterior is on top. Scale bars 100 μm (**A**) and 25 μm (**A’**). **B**-**B’**-Confocal projections focusing on endothelial cells (in green) from the dorsal aorta (DA) and the cardinal vein (CV) at 24 hpf. Red and blue arrows point to Prmt5+ cells (in magenta) from the DA and the CV, respectively. Red and blue lines represent DA and CV diameters, respectively. Scale bar 50 μm. **C**-**C’**-Confocal projections focusing on sprouting ISVs (in green) at 24 hpf. Light blue and yellow arrows point to tip and stalk cell, respectively. **D**-Schematic representation of the trunk vasculature with ISVs sprouting from the DA. The tip cell leads the cell migration and the stalk cell maintains the connection with the DA. **E**-**F**-Confocal projections of transgenic *Tg(fli1a:GFP)^y1^* wild type (**E**) and *prmt5* mutant (**F**) embryos at 28 hpf. Red rectangles delimit where DA close ups were made. White rectangles delimit the higher magnification (x2) of the DA with red lines indicating the dorsal aorta diameters. White arrows indicate the connection point between two ISVs to form the Dorsal Longitudinal Anastomotic Vessel (DLAV). Scale bar 100 μm. **G**-**H**-Confocal projections of control morphant (**G**) and *prmt5* morphant (**H**) transgenic *Tg(TP1bglob:VenusPEST)^s940^* embryos labelling cells from the DA at 28 hpf. Yellow lines delimit the measured area occupied by the DA. Scale bar 25 μm **I**-Average area occupied by the DA in μm^2^ in control and *prmt5* morpholino injected embryos from 2 independent experiments with at least 8 animals per condition. T-test was performed. **J**-**K**-Average number of endothelial cells per intersegmental vessel (**J**) and average ISV length in μm (**K**) in control and in *prmt5* mutant embryos from 3 independent experiments with at least 3 animals per condition. T-test and Mann Whitney test were performed, respectively. ** P<0.01, ***P<0.001.

To get a better insight into the impact of Prmt5 on the dynamics of vascular system formation, we performed time-lapse analyses in control and *prmt5* morphant embryos. Time-lapse confocal movies were carried out from 28 hpf to 38 hpf to follow the elongation of ISVs to the formation of an effective lumen. As compared to control morphants, *prmt5* morphants showed an impaired formation of ISV lumen and DLAV. Indeed, *in prmt5* morphants tip cells failed to stay connected to the stalk cells and to contact other tip cells to allow the formation of the DLAV (Fig. 3A-B). Moreover, supernumerary connections were detected in the context of *prmt5*-loss of function (Fig. 3B). Altogether, these data suggest a central role for Prmt5 in vascular morphogenesis.

**Figure 3:**
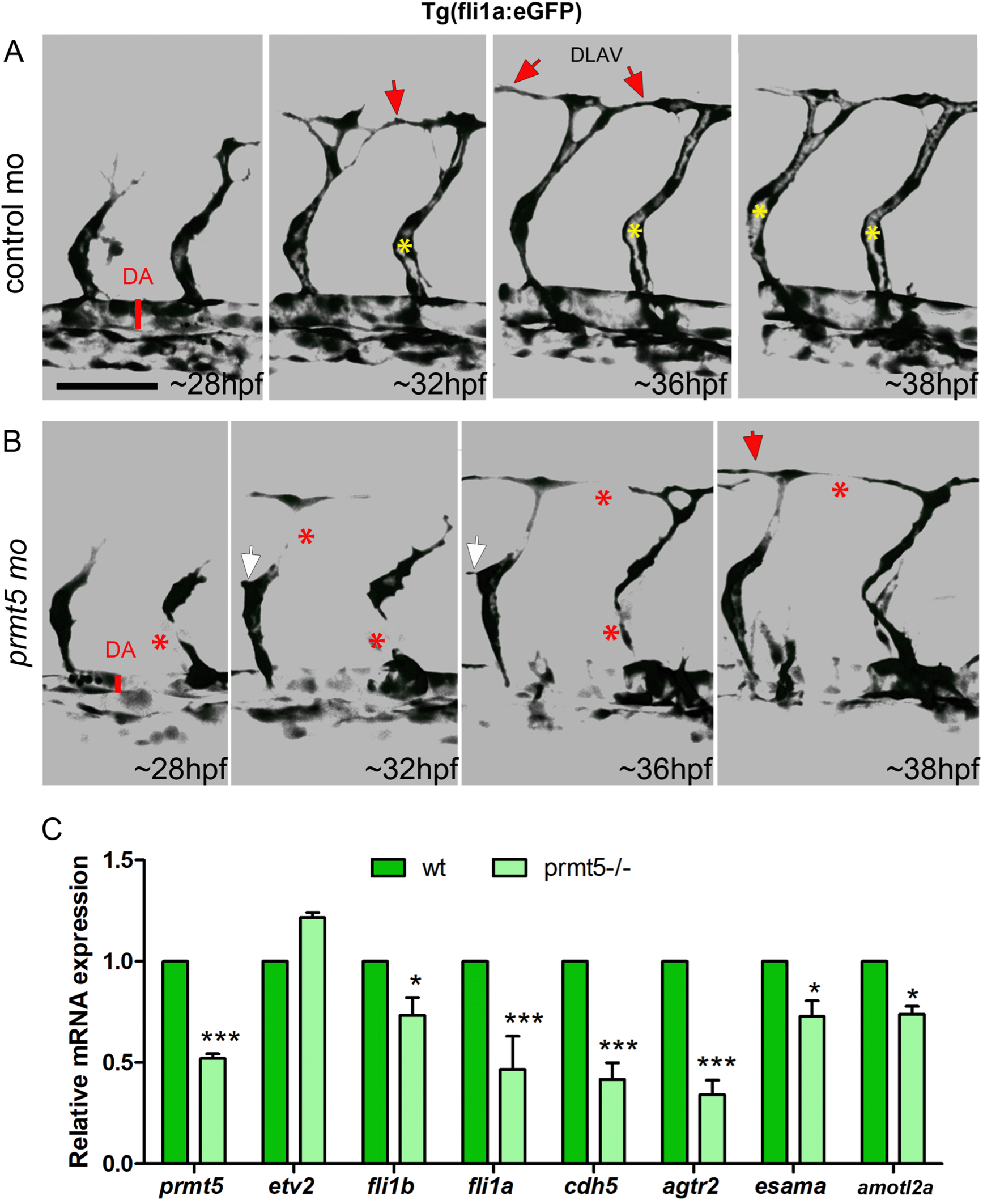
Prmt5 is required for vascular morphogenesis. **A**-**B**-Still images from movies of control (**A**) and *prmt5* morphant (**B**) Tg(fli1a:GFP)^y1^ transgenic embryos from 28 to 38 hpf. Red asterisks label missing connections between tip and stalk cells as well as missing connections between tip cells that should lead to DLAV formation. Red arrows point to connecting ISVs leading to DLAV formation. White arrows indicate supernumerary sprouts. Yellow asterisks label the lumen of ISVs. Scale bar 50 μm. **C**-Relative mRNA expressions of the indicated transcripts were determined by RT-qPCR in 28 hpf wild type and *prmt5* mutant embryos, from 3 independent experiments with at least 6 animals per condition. Two-way ANOVA was performed. * P<0.05, ***P<0.001.

The master gene regulator ETV2, ETS transcription factors and adhesion proteins have been shown to be involved in blood vessel formation (Craig et al., 2015; Hultin et al., 2014; Pham et al., 2007; Sauteur et al., 2017; Sauteur et al., 2014). Analyzing single cell RNA-sequencing data from Wagner et al. (Wagner et al., 2018), allowed us to determine that *prmt5* is expressed in endothelial cells at 10 hpf (like *etv2* and *fli1a*) and that its expression decreases in later stages, when the expression of *fli1b*, *cdh5*, *agtr2*, *esama*, and *amotl2a* starts to increase (Fig. S3). To test whether Prmt5 could regulate the expression of these genes, we performed RT-qPCR experiments on mutant embryos and on their wild type counterparts. While we found that *etv2* expression was not affected, the expression of ETS transcription factors (*fli1a, fli1b*) and adhesion proteins *(cdh5*, *agtr2*, *esama* and *amotl2a),* all putative ETV2 target genes (Liu et al., 2015b; Wong et al., 2009), was significantly reduced in *prmt5* mutant (Fig. 3C). Of note, we also detected a reduction of *fli1a* and *cdh5* expression in *prmt5* mutant by *in situ* hybridization (Fig. S4). As *etv2* expression was unaffected by the loss of *prmt5* but its targets were down-regulated, it is tempting to speculate that Prmt5 could modulate ETV2 activity at post-translational level.

### Prmt5 methyltransferase activity is not required for vascular morphogenesis

That Prmt5 modulates gene expression by methylating a variety of proteins including histones but also transcription (co)factors led us to test whether Prmt5 methyltransferase activity was required for vascular morphogenesis and lymphoid progenitor formation. To this end, *prmt5* mutant or morphant embryos were injected with wild type *human prmt5* mRNA (*hprmt5WT*) or with a catalytic mutant form of this mRNA (*hprmt5MUT*) (Pal et al., 2003). In mice, the expansion of lymphoid progenitor relies on Prmt5 methyltransferase activity (Liu et al., 2015a). Consistent with this, *hprmt5WT* but not *prmt5MU*T mRNA, was able to restore normal lymphoid progenitor expansion in *prmt5* morphant embryos (Fig. S1 C-G). This underscores the conserved requirement of PRMT5 methyltransferase activity for lymphoid progenitor formation in human and zebrafish. We then tested whether the same was true for ISV elongation and the expression of *etv2* target genes. Surprisingly, we found that both mRNAs were able to restore ISV elongation, albeit to a slightly different extend, as indicated by the average ISV length in injected mutant embryos as compared to non-injected mutants (Fig. 4A-E). Indeed, we observed that the average length of ISVs in *hprmt5WT*-injected mutants was even longer than intersegmental vessels of wild type embryos, while the average length in *hprmt5MUT* injected mutants was significantly superior to non-injected mutants but shorter than control embryos (Fig. 4 E). Interestingly, no difference could be seen in the cell number per ISV in the different contexts (Fig. 4F) thus ruling out the possibility that Prmt5 regulates cell proliferation at the ISV. Finally, RT-qPCR experiments revealed that both *hprmt5WT* and *hprmt5MUT* mRNAs were able to restore the expression of *etv2* target genes, except for *fli1a* whose expression was only rescued by *hprmt5WT* (Fig. 4G). In sum, these results indicate that Prmt5 methyltranferase activity is largely dispensable for its function in blood vessel formation.

**Figure 4:**
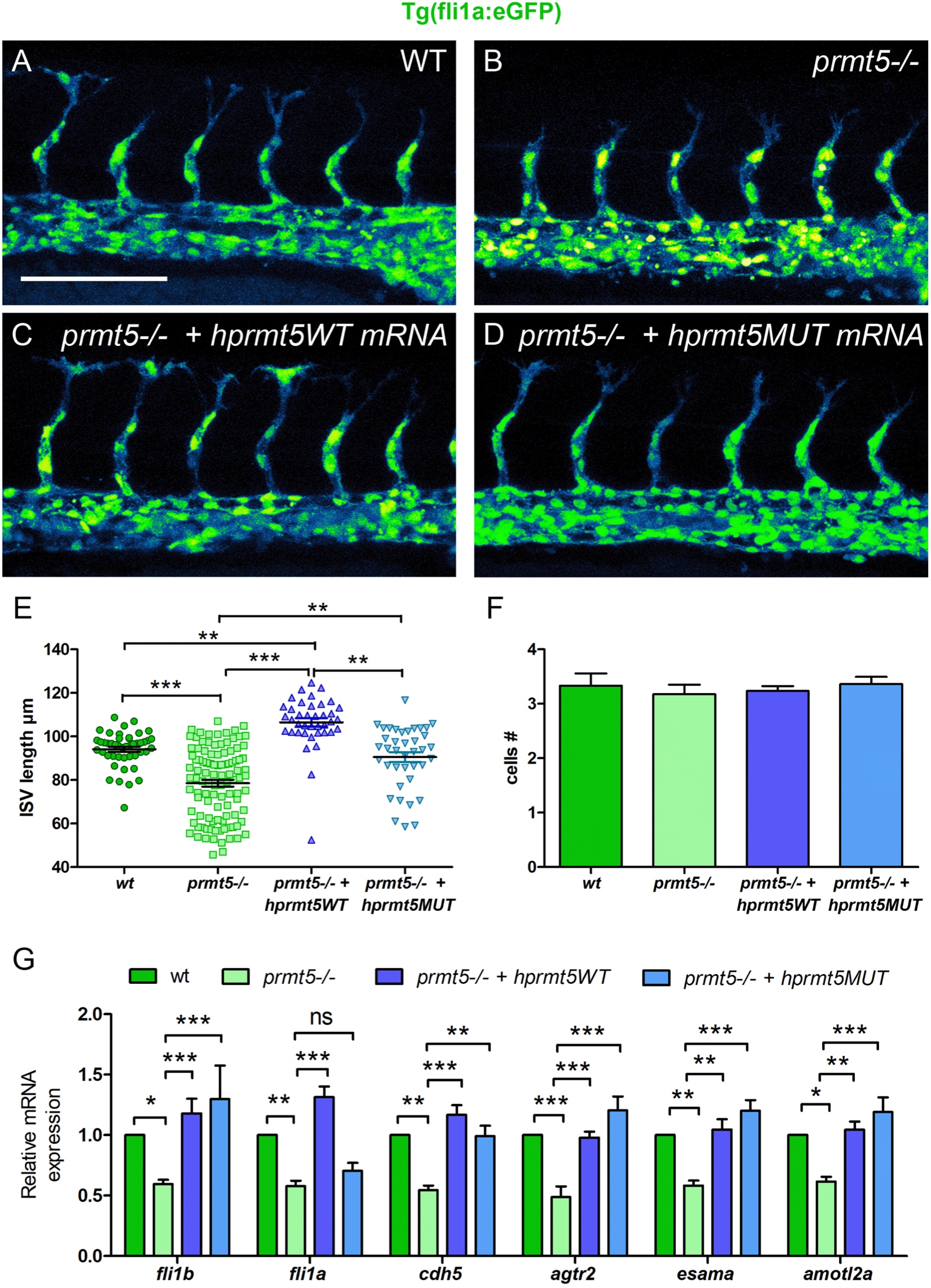
Prmt5 methyltransferase activity is dispensable for vascular morphogenesis. **A**-**D**-Confocal projections of transgenic *Tg*(*fli1a:GFP*)^y1^ embryos at 28 hpf. Wild type embryo is on the top left panel (**A**), *prmt5* mutant embryos were not injected (**B**) or injected with either *hprmt5WT* mRNA (**C**) or the mutant form *hprmt5MUT* mRNA (**D**). Scale bar 100 μm. **E**-**F**-Average ISVs length in μm (**E**) and average number of endothelial cells per ISVs (**F**) for wild type, *prmt5* mutant embryos not injected or injected with *hprmt5WT* mRNA, or *hprmt5 MUT* mRNA, from 3 independent experiments with at least 3 animals per condition. Kruskal-Wallis test (**E**) and One-way ANOVA (**F**) were performed. ** P<0.01, *** P<0.001. **G**-Relative mRNA expressions were determined by RT-qPCR on 28 hpf wild type and *prmt5* mutant embryos injected by either *hprmt5WT or hprmt5MUT* mRNAs, from 2 independent experiments with at least 6 animals per condition. Two-way ANOVA was performed. * P<0.05.

### Prmt5 might help to shape correct chromatin conformation in endothelial cells

As Prmt5 methyltransferase activity seems to be not required for gene expression regulation in vascular morphogenesis, we speculated that Prmt5 could act as a scaffold protein in complexes mediating transcription and chromatin looping. Indeed, Prmt5 has been proposed to promote enhancer-promoter looping at the PPARγ2 locus and more broadly to facilitate chromatin connection in adipocytes, *via* the recruitments of Mediator subunit MED1 and SWI/SNF chromatin remodeling complex subunit Brg1 ATPase (LeBlanc et al., 2016). Thus, we decided to inspect the chromatin architecture of the flanking region of identified Prmt5-regulated genes using ATAC-seq data from zebrafish endothelial cells that we previously generated (Quillien et al., 2017). Doing so, we found that putative enhancers are on average distant of 16 kb from the transcriptional start site (TSS) (Table S1, Figure S5), indicating that their expression could rely on proper chromatin looping. To further characterize these specific *cis* regulatory regions, we turned into the mouse model and analyzed the ChIP-seq data of Etv2 and Prmt5-dependent H4R3 di-methylation to determine whether Prmt5 target genes identified in our study were conserved in mouse (Girardot et al., 2014; Liu et al., 2015a). We found that Etv2 is recruited to the cis regulatory element of *amotl2*, *cdh5* and *fli1* (Table 1) and that its binding was associated with the presence of H4R3me2 for some of them, suggesting that ETV2 and Prmt5 can be recruited on the same regions in mouse. In order to gain further insight into the potential role of Prmt5 in supporting proper chromatin conformation in endothelial cells, we analyzed and compared the expression of *Gal4* reporter genes in an endogenous (Fig. 5A) and in an artificial chromatin context (Fig. 5E). The first construction used consists in the transgenic line *TgBAC(cdh5:GAL4FF);Tg(UAS:GFP)* that contains the sequence of an optimized version of Gal4VP16 (*GAL4FF*) inserted at the TSS of *cdh5* gene between *cdh5* promoter region (P) and a putative enhancer (E) distant of ~20kb as defined by the presence of two ATAC-seq positive regions (Table S1, Fig. 5A, Fig. S5) (Bussmann and Schulte-Merker, 2011; Quillien et al., 2017). Therefore, in double transgenic individuals, the level of GFP fluorescence intensity correlates with endogenous *cdh5* expression. In addition, we generated a transgenic line where the *cdh5* promoter and putative enhancer were cloned next to each other, both upstream of the *Gal4VP16* coding sequence (Fig. 5E). In double transgenic embryos *Tg(cdh5:Gal4VP16); Tg(UAS:KAEDE),* the fluorescence intensity of the protein KAEDE is an artificial read out of *cdh5* transcription for which chromatin looping is not required. Comparing the level of fluorescence intensity in *TgBAC(cdh5:GAL4FF);Tg(UAS:GFP)* transgenic line in control condition and in the context of *prmt5* knock down, we observed a strong reduction of GFP fluorescence intensity in *prmt5* morphants (Fig. 5B-D), indicating that Prmt5 is required for *cdh5* expression in an endogenous context. In double transgenic embryos *Tg(cdh5:Gal4VP16); Tg(UAS:KAEDE)*, the fluorescent protein KAEDE was expressed in blood vessels (Fig. 5F), validating that the putative enhancer and the promotor region of *cdh5* are sufficient to drive gene expression in endothelial cells. However, in this artificial context, *prmt5* morpholino injection had no effect on the level of KAEDE fluorescence intensity as compared to control morphants (Fig. 5F-H). This result suggests that in this particular context *i.e.* when chromatin looping between enhancer and promoter was not needed, Prmt5 was not required either for gene expression. This finding supports the idea that Prmt5 plays a role in the formation of the correct 3D environment for endothelial genes expression. Finally, rescue experiments were performed by injecting either wild type or a catalytic mutant of human *prmt5* mRNA to determine whether Prmt5 methyltransferase activity was required for the transcriptional control of *cdh5* expression in the endogenous context. We found that both wild type and mutant *hprmt5* mRNAs restored GFP fluorescence intensity in *prmt5* morphants as compared to control embryos (Fig. 5B-D, I-J). Collectively, these data indicate that the transcriptional control of *cdh5* is independent of Prmt5 methyltransferase activity and could rather rely on a role of Prmt5 as a scaffold protein to provide a proper chromatin conformation context.

**Figure 5:**
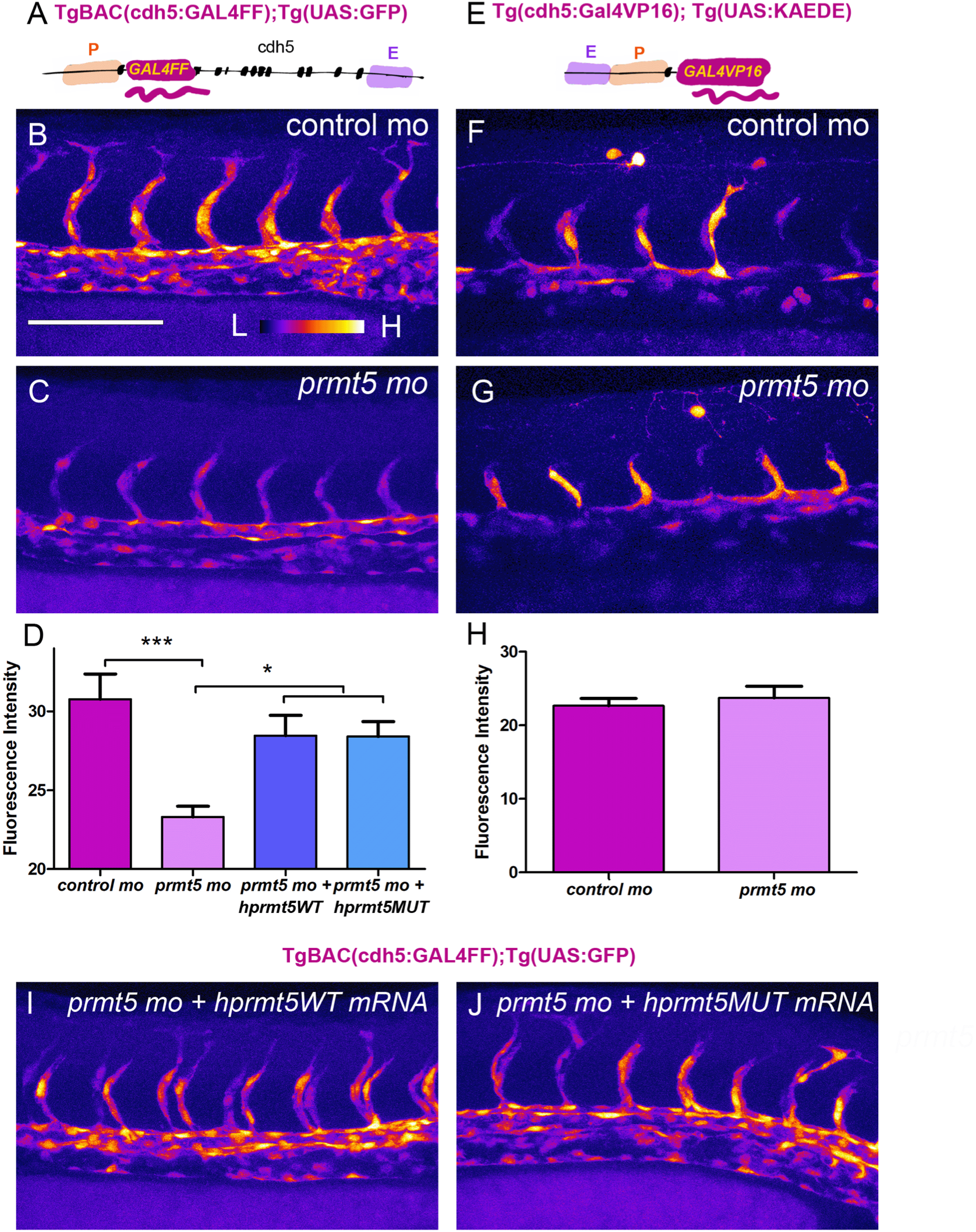
Prmt5 promotes chromatin looping. **A**-Schematic representation of the transgene *TgBAC(cdh5:GAL4FF)* containing two putative cis-regulatory elements, a promotor region (P) and an enhancer (E), separated by ~20kb with the *GAL4FF* reporter gene inserted at the TSS of *cdh5.* **B**, **C**, **I**, **J**-Confocal projections of transgenic *TgBAC(cdh5:GAL4FF);*Tg(*UAS:GFP*) embryos at 28 hpf. Control morphant is on the top left panel (**B**), *prmt5* morphant embryos were not injected (**C**) or injected by either *hprmt5WT* mRNA (**I**) or the catalytic mutant form *hprmt5MUT* (**J**) mRNA. The fluorescent intensity is colored-coded, from the Low intensity (L) in black to High intensity (H) in white (intensity scale as in panel **B**). Scale bar 100 μm. **D**-Average GFP fluorescence intensity per confocal projection for control, *prmt5* morphant embryos injected by *hprmt5WT* mRNA, or *hprmt5 MUT* mRNA or not injected, from 3 independent experiments with at least 3 animals per condition. One-way ANOVA was performed. *P<0.05, ***P<0.001. **E**-Schematic representation of the transgene Tg(*cdh5:GAL4VP16*) containing the two putative cis-regulatory elements next to each other (E and P), upstream of *GAL4VP16* reporter gene. **F**-**G**-Confocal projection of transgenic Tg(*cdh5:GAL4VP16*);Tg(*UAS:KAEDE*) embryos at 26 hpf injected with either a control morpholino (**F**) or *a prmt5* morpholino (**G**). The fluorescence intensity is color-coded, from the Low intensity (L) in black to High intensity (H) in white (intensity scale in panel **B**). **H**-Average KAEDE fluorescence intensity for control and for *prmt5* morphant embryos, from 3 independent experiments with at least 5 animals per condition. T-test was performed.

**Table 1:**
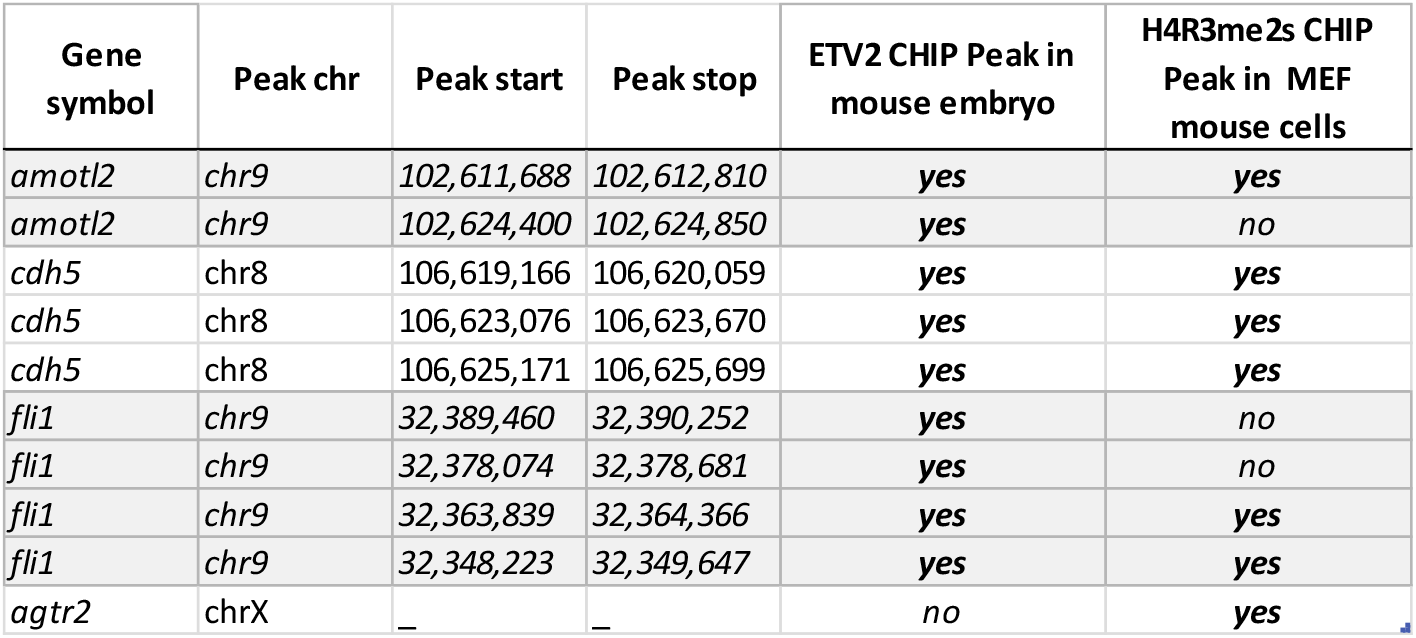
Chromatin profile of mouse orthologous genes of prmt5 identified target genes. List of Etv2 and H4R3me2s peaks identified by CHIP-seq in mouse embryos and in MEF mouse cells, respectively (Liu et al., 2015a; Girardot et al., 2014).

## DISCUSSION

Here we have demonstrated a role for Prmt5 in both hematopoiesis and blood vessel formation in zebrafish. Our results suggest that Prmt5 promotes vascular morphogenesis through the transcriptional control of ETS transcription factor and adhesion proteins in endothelial cells. Intriguingly, we have shown that the methyltransferase activity of Prmt5 was not absolutely required to regulate gene expression, leading us to propose a role of scaffold protein for Prmt5 to facilitate chromatin looping formation in endothelial cells.

We found that, similarly as in mouse (Liu et al., 2015a), Prmt5 plays an important role in zebrafish hematopoiesis by controlling HSCs emergence and HSPCs expansion. We also described for the first time the involvement of Prmt5 in vascular morphogenesis by regulating the expression of known genes that control this process (adhesion proteins or transcription factors). Actually, *prmt5* loss of function partially phenocopied loss of function of these genes. Indeed, it was shown that knocking down individual ETS proteins had limited effect on sprout formation, while the combination of morpholinos against both *fli1a, fli1b,* and *ets1* led to a decreased number of vessel sprouts at 24 hpf but to a normal trunk vasculature at 48 hpf (Pham et al., 2007). Moreover, *amolt2a* knock down in zebrafish led to a reduced diameter of the DA in a similar way as we found in the context of *prmt5* loss of function (Hultin et al., 2014). Furthermore, disconnected stalk and tip cells and delayed formation of the DLAV formation that we observed in *prmt5* mutant phenocopies loss of function of both *cdh5* and *esama* published in previous studies (Sauteur et al., 2017; Sauteur et al., 2014). However, the loss of function of *cdh5* had no effect on HSCs emergence or HSPCs expansion (Anderson et al., 2015), suggesting that Prmt5 might act on different set of genes in endothelial cells and in emerging HSCs. In agreement with this hypothesis, Tan et al. have proposed that Prmt5 is playing a critical role in HSC quiescence through the splicing of genes involved in DNA repair (Tan et al., 2019). Of note, this study showed that Prmt5 methyltransferase activity was required for controlling HSC quiescence, in agreement with our findings in the present work. In contrast, our data suggest that the methyltransferase activity of Prmt5 is dispensable in endothelial cells, reinforcing the idea that Prmt5 regulates transcription by different mechanisms in these two processes (Fig. 6).

**Figure 6:**
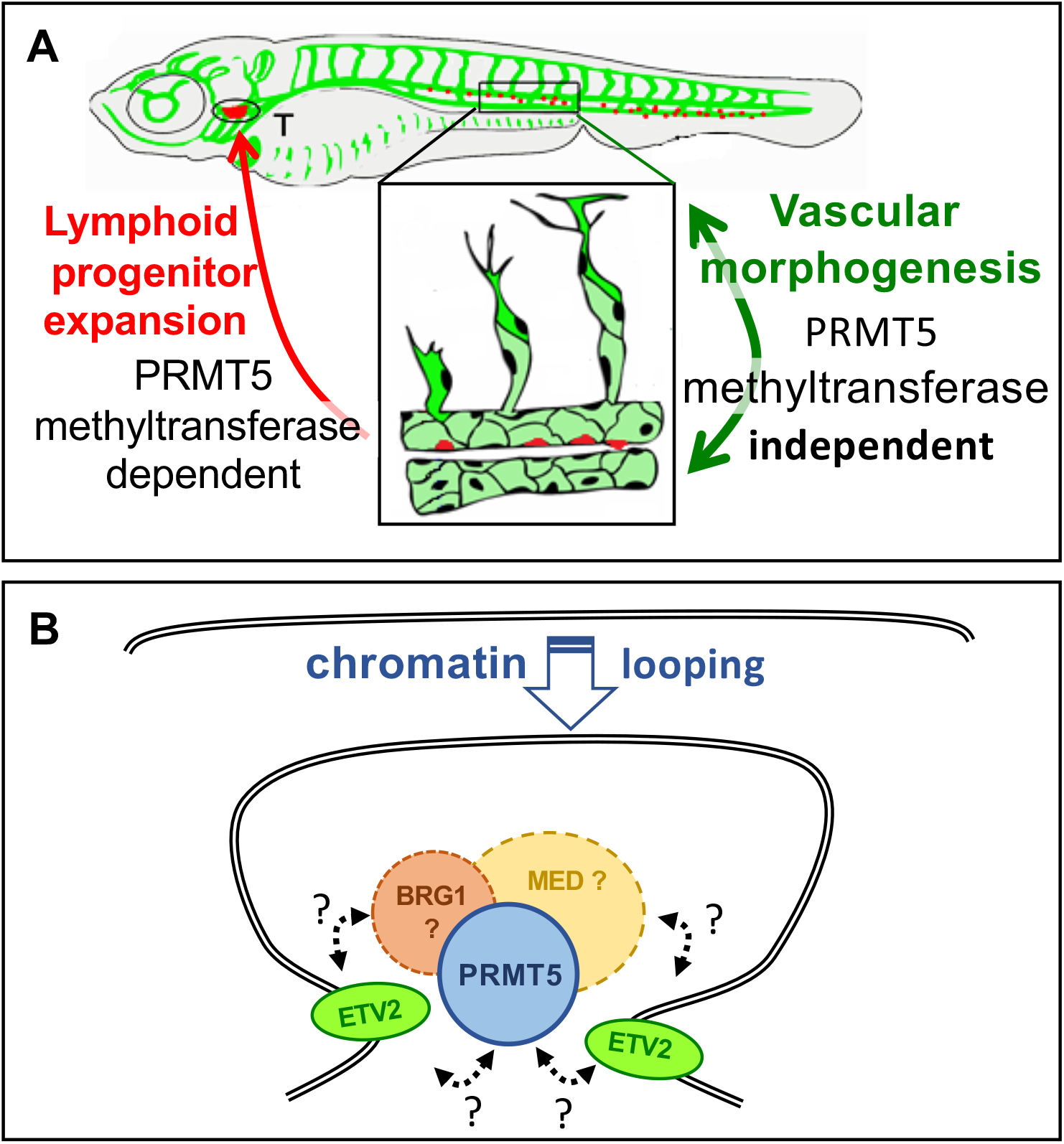
**A-**Schematic representation of two distinct roles of Prmt5 during the formation of hematopoietic lineage development and blood vessels, relying or not on its methyltransferase activity, respectively. **B**-Proposed model to depict the function of Prmt5 in zebrafish endothelial cells. The transcription factor ETV2 recruited to promoters and enhancers of endothelial specific genes, could favor the recruitment of a complex including Prmt5, Brg1 and the mediator complex to help the formation of chromatin loping and thus facilitate the transcription of specific endothelial genes. Dashed lines indicate potential interactions or plausible recruitments of Brg1 and/or the mediator complex.

Prmt5 has been shown to facilitate ATP-dependent chromatin remodeling to promote gene expression in skeletal muscles and during adipocyte differentiation (Dacwag et al., 2009; LeBlanc et al., 2012; LeBlanc et al., 2016; Pal et al., 2003). Here, we propose that Prmt5 could also be essential for proper chromatin looping in endothelial cells. Our data suggest that Prmt5 influences gene expression only in an endogenous context where chromatin looping is required (*e.g*. *chd5* and *TgBAC(cdh5:GAL4FF))*, while it is dispensable for gene expression when enhancer and promotor regions are artificially associated (*e.g. Tg(cdh5:Gal4VP16))* or close by (*e.g*. *fli1a*). This implies that Prmt5 could interact with Brg1 ATPase of SWI/SNF chromatin remodeling complex and with the Mediator complex in endothelial cells as it does in muscle cells and adipocytes. Consistent with this hypothesis, *brg1* mutant mouse embryos display an anemia coupled to vascular defects in the yolk sac, characterized by thin vessels and supernumerary sprouts (Griffin et al., 2008), which is reminiscent to our present findings in zebrafish *prmt5* mutant. Interestingly, it has been proposed that the mediator complex regulates endothelial cell differentiation (Napoli et al., 2019). Moreover, our analyses of the published single cell expression data (Wagner et al., 2018) indicate that, similarly to *prmt5*, the expression of *smarc4a/brg1* and *med12* in zebrafish endothelial cells is detected as early as 10 hpf and decreases in subsequent stages. It is thus tempting to speculate that Prmt5, Brg1 and the Mediator could act together to regulate chromatin organization in endothelial cells (Fig. 6).

ChIP-seq data available in mouse revealed that some flanking regions of orthologues of identified Prmt5 target genes are bound by ETV2 and present histone marks associated with the recruitment of Prmt5. In zebrafish, both *prmt5* and *etv2* genes are expressed at early stage in endothelial cells, and Etv2 binding motif is enriched in *cis*-regulatory regions identified by ATAC-seq experiment (Quillien et al., 2017). In addition, zebrafish mutant for *prmt5* from our study and a mutant for the master regulator *etv2* shared similarities in their phenotypes displaying abnormal vasculature at 48 hpf characterized by a lack of lumen formation, a lack of vessel extension and aberrant connections (Craig et al., 2015; Pham et al., 2007). Here, we proposed that Etv2 could be involved in the recruitments of Prmt5 to *cis* regulatory regions of endothelial genes. Another crucial player of blood vessel formation is the transcription factor Npas4l, which is expressed during late gastrulation and regulates *etv2* expression (Marass et al., 2019). Npas4l ChIP-seq data and ATAC-seq data from *npas4l* mutant also revealed the binding of this transcription factor to a certain number of cis-regulatory regions of Prmt5 target genes identified in the present work. In light of these findings, we speculate that Npas4l could contribute to the recruitment of Prmt5 to endothelial genes (AQ and LV, unpublished data). Even though technically highly challenging at the present time, ChIP-seq against Prmt5 or any known Prmt5 substrates in endothelial cells in zebrafish combined with the corresponding RNA-seq/ATAC-seq experiments in wild type or mutant condition for Prmt5 could help to validate our model and identify all Prmt5 putative target genes.

The presence of Prmt5 and Brg1 at promotor regions of the *PPARγ2* locus or of *myogenin* was associated with dimethylated H3R8 (histone 3 arginine 8) (Dacwag et al., 2009; LeBlanc et al., 2012). Interestingly, *prmt5* knock down led to a reduction of both histone methylation and chromatin looping formation (Dacwag et al., 2009; LeBlanc et al., 2012; LeBlanc et al., 2016). *In vitro*, the addition of Prmt5 to Brg1-immunopurified complexes enhanced histone methylation, while the addition of a catalytic dead version of Prmt5 did not (Pal et al., 2003). Altogether these data suggest that wild type Prmt5, when recruited to target gene promoter regions, acts most likely by dimethylating histone proteins. However, these studies did not assess the ability of Prmt5 to facilitate chromatin looping independently (or not) of its methyltransferase activity. Our data suggest that chromatin looping favored by Prmt5 does not necessarily require its methyltransferase activity. Indeed, rescue experiments demonstrated that Prmt5 was able to restore gene expression independently of its enzymatic activity, with the exception of *fli1a* expression. Since *fli1a* putative enhancer is located only at 700 pb from the promoter region, chromatin looping might not be required for *fli1a* expression and Prmt5 might essentially act here through its methyltransferase activity. Hence, depending on the context and the target genes considered, Prmt5 could modulate gene expression in endothelial cell through promotion of chromatin interaction and/or *via* histones/proteins modification. Finally, we can consider that other proteins of the PRMT family could also regulate endothelial gene expression, as some PRMT members are also expressed in zebrafish endothelial cells (AQ and LV, unpublished data). For instance, ChIP-seq data in chicken erythrocytes suggest that both Prmt5 and Prmt1 are recruited to the same cis-regulatory regions with Prmt1 permitting the recruitments of CBP/p300 to acetylate histones (Beacon et al., 2020). Hence, analyses of the role(s) of other PRMT family members in endothelial cells would help to better understand the cross-talks between these enzymes. Besides their function during normal development, it has been shown in a zebrafish xenotransplantation model that Etv2 and Fli1b are required for tumor angiogenesis, suggesting that inhibition of these ETS factors may present a novel strategy to inhibit tumor angiogenesis and reduce tumor growth (Baltrunaite et al., 2017). We found that Prmt5 activates ETV2 target gene expression, and Prmt5 has been proposed as a therapeutic target in many diseases, including cancer (Shailesh et al., 2018). Several Prmt5 inhibitors have been discovered in the past decade and some have been tested in clinical trials for the treatment of tumors (reviewed in (Wang et al., 2018)). However, the vast majority, if not all, compounds discovered and validated so far inhibit Prmt5 enzymatic activity (Lin and Luengo, 2019). Yet, we show here that Prmt5 acts at least in part, independently of its methyltransferase activity to regulate vascular morphogenesis. Hence, our data shed light on a mechanism of action of Prmt5 that will be insensitive to the afore mentioned enzymatic inhibitors and thus calls forth the design of alternative drugs *i.e*. specific inhibitors of the interaction between Prmt5 and Etv2 in this context. In conclusion, our study highlights different modes of regulation of gene expression by Prmt5 in endothelial cells and strengthens the importance of its enzymatic-independent function in chromatin looping. This non-canonical function of Prmt5 may have a more pervasive role than previously thought in physiological conditions *i.e.* during development but also in pathological situations such as in tumor angiogenesis and this aspect certainly deserves more attention in the future.

## MATERIALS AND METHODS

### Zebrafish care and maintenance

Embryos were raised and staged according to standard protocols and the Recommended Guidelines for Zebrafish Husbandry Conditions (Alestrom et al., 2019; Kimmel et al., 1995). The establishment and characterization of Tg(gata2b:Gal4;UAS:lifeactGFP), Tg(fli1a:eGFP), Tg(TP1bglob:VenusPEST)s940, TgBAC(cdh5:GAL4FF);Tg(UAS:GFP), Tg(UAS:KAEDE) have been described elsewhere (Bussmann and Schulte-Merker, 2011; Butko et al., 2015; Hatta et al., 2006; Lawson and Weinstein, 2002b; Ninov et al., 2012). Lines generated in this study are described below. Embryos were fixed overnight at 4°C in BT-FIX, after which they were immediately processed or dehydrated and stored at −20°C until use.

### Ethics statement

Fish were handled in a facility certified by the French Ministry of Agriculture (approval number A3155510). The project has received an agreement number APAFIS#7124-20161 00517263944 v3. Anesthesia and euthanasia procedures were performed in Tricaine Methanesulfonate (MS222) solutions as recommended for zebrafish (0.16 mg/ml for anesthesia, 0.30 mg/ml for euthanasia). All efforts were made to minimize the number of animals used and their suffering, in accordance with the guidelines from the European directive on the protection of animals used for scientific purposes (2010/63/UE) and the guiding principles from the French Decret 2013–118.

### Plasmid construction

To construct the transgene Tg(*cdh5:GAL4VP16*), we cloned the putative *cdh5* promoter (*cdh5*P) and enhancer (*cdh5*E) elements into pme_mcs and p5E_GGWDest+ (Addgene #49319) (Kirchmaier et al., 2013; Kwan et al., 2007) using XhoI, EcoRI and BsaI to give pme_*cdh5*P and p5E_*cdh5*E, respectively. The Gal4VP16 sequence from pme_Gal4VP16 (Kwan et al., 2007) was then introduced downstream of *cdh5*P into pme_*cdh5*P using BamH1 and SpeI. A multisite LR recombination reaction (Gateway LR Clonase II Enzyme mix, Invitrogen) was then performed using p5E_*cdh5*E, pme_*cdh5*P:Gal4VP16, with pminTol-R4-R2pA to give pminTol-*cdh5*E-*cdh5*P: Gal4VP16. Oligonucleotide sequences are listed in Table S2.

### Generation of *prmt5*^−/−^ mutants by CRISPR/cas9

The guide RNA (gRNA) was designed using CHOPCHOP CRISPR Design website (Montague et al., 2014). The designed oligos were annealed and ligated into the gRNA plasmid pDR274 after digestion of the plasmid with BsaI (NEB). The gRNA was prepared *in vitro* using the MEGAshortscript T7 transcription kit (Ambion) after linearizing the plasmid with DraI (NEB) (Talbot and Amacher, 2014) before being purified using illustra MicroSpin G-50 Columns (GE Healthcare). 1 nL of a solution containing 10μM EnGen Cas9 NLS (NEB) and 100 ng/μl of gRNA was injected at the one-cell stage. WT, heterozygous, and homozygous *prmt5* animals were identified by PCR. Oligonucleotide sequences are listed in Table S2.

### Microinjections

The Tg(*cdh5:GAL4VP16*);Tg(*UAS:KAEDE*) line was generated using pminTol-*cdh5*E-*cdh5*P: Gal4VP16 by Tol2 transposition as described previously (Covassin et al., 2009). Control and *prmt5* morpholino oligonucleotides (MOs) were described previously (Batut et al., 2011). Embryos from in-crosses of the indicated heterozygous carriers or wild-type adults were injected at the one cell stage with 6 ng of MO. pBluescript II KS+ hPRMT5 WT and pBluescript II KS+ hPRMT5 Mutant (Pal et al., 2003) were linearized by EcoRI (NEB) and transcribed by T7 (Promega). 200 pg *hprmt5WT* mRNA, or *hprmt5 MUT* mRNA were injected at one cell stage.

### RNA extraction, Reverse transcription and real-time PCR

Embryos were dissected at the indicated stage after addition of Tricaine Methanesulfonate. Genomic DNA was extracted from dissected embryo heads to identify their genotype and the corresponding dissected tails were conserved in TRIzol Reagent at −20°C. After identification of wild type and mutant embryos, total RNAs from at least 6 identified tails were extracted following manufacturer’s instructions (Invitrogen). Total RNAs were converted into cDNA using Prime Script cDNA Synthesis Kit (Takara) with Oligo(dT) and random hexamer primers for 15 min at 37 °C according to manufacturer’s instructions. cDNAs were then diluted 20-fold and quantified by qPCR using SsoFast Evagreen Supermix (Bio-rad) and specific primers. Data were acquired on CFX96 Real-Time PCR detection System (Bio-rad). Samples were analyzed in triplicates and the expression level was calculated relative to zebrafish housekeeping gene *EF1α*. Oligonucleotide sequences are listed in Table S2.

### Live imaging

For the transgenic lines *TgBAC(cdh5:GAL4FF);*Tg(*UAS:GFP*) and Tg(*cdh5:GAL4VP16*);Tg(*UAS:KAEDE*), embryos were placed in 1.5% low melt agarose with Tricaine on a glass-bottomed culture dish filled with egg water. Images were acquired using the confocal microscope TCS SP8 (Leica Microsystems) with an L 25 × /0.95 W FLUOSTAR VIZIR objective (zoom X1.25) using the scanner resonant mode. Confocal stacks were acquired every 10 min from 28 to 38 hpf to generate movies.

### Immunostaining and *in situ* hybridization

After fixation or rehydratation, embryos were washed twice with Phosphate Buffered Saline/1% Triton X-100 (PBST), permeabilized with PBST/0.5% Trypsin for 30 sec and washed twice again with PBST. After blocking with PBST/10% Fetal Calf Serum (FCS)/1% bovine serum albumin (BSA) (hereafter termed ‘blocking solution’) for at least 1 h, embryos were incubated with antibodies directed against either GFP (Torrey Pine, Biolabs), or Prmt5 (Upstate #07405), in blocking solution overnight at 4 °C followed by 5 washing steps with PBST. Embryos were then incubated with the appropriate Alexa Fluor-conjugated secondary antibodies (Molecular Probes) for at least 2 h at room temperature and washed three times. Nuclei were then stained with TO-PRO3 (Molecular Probes) and washed twice with PBST. Embryos were dissected, flat-mounted in glycerol and images were recorded on a confocal microscope as above. Fluorescent *in situ* hybridization was carried out as previously described (Quillien et al., 2014).

### Image processing and measurements

Confocal images and stacks were either analyzed with ImageJ software or LAS X. Nuclei of ISV cells and gata2b+ cells were counted using the Multipoint tool of ImageJ. ISV lengths were measured by drawing a line between the base and the tip of ISV on ImageJ. Contours of the Dorsal Aorta were drawn using the Freehand Selection Tool with a digital pen and the area was then measured. Fluorescence intensity corresponded to the measurement of average gray value for each entire image.

### Statistical analysis

Statistical comparisons of datasets were performed using GraphPad Prism software. For each dataset, we tested the assumption of normality with D’Agostino-Pearson tests and accordingly, unpaired t-test, Mann-Whitney test, One-way ANOVA, two-way ANOVA or Kruskal-Wallis test were used to compare dataset; means (± SEM) are indicated as horizontal bars on dot plots. The test used as well as the number of independent experiments performed and the minimal number of biological replicates are indicated in each figure legend.

### Bioinformatic analysis

Published single cell data from total embryos at 10hpf, 14hpf, 18hpf and 24hpf (Wagner et al., 2018) were analyzed using the R package Seurat (Butler et al., 2018; Stuart et al., 2019). After data clustering, clusters of endothelial cells from each stage were identified by the expression of several endothelial specific genes (*etv2*, *fli1a*, …). Then, we examined the level of expression and the percentage of cells expressing our gene of interest at each developmental stage. ATAC-seq data (Marass et al., 2019; Quillien et al., 2017) and Chip-seq data (Girardot et al., 2014; Liu et al., 2015a) were inspected using the Galaxy platform (Afgan et al., 2018).

## ACKNOWLEDGEMENTS

We would like to thank the Blader lab for helpful discussions and support; Dr. Waltzer, Dr. Cau, Dr. Male and Dr. Navajas Acedo for critical reading of the manuscript; A. Laire for excellent zebrafish care; B. Ronsin and S. Bosch from the Toulouse Rio Imaging (TRI) platform; M. Aguirrebengoa from the BigA Core Facility, CBI Toulouse; Dr. Herbomel for the Zebrafish transgenic line *Tg(gata2b:Gal4;UAS:lifeactGFP).* This work was supported by a grant from the Fondation de l’Association pour la Recherche contre le Cancer (Fondation ARC) and from the Association Française contre les Myopathies (AFM) to LV. AQ was supported by a fellowship from the Fondation ARC.

## COMPETING INTEREST

The authors declare no competing interests.

---

**Figure S1:**
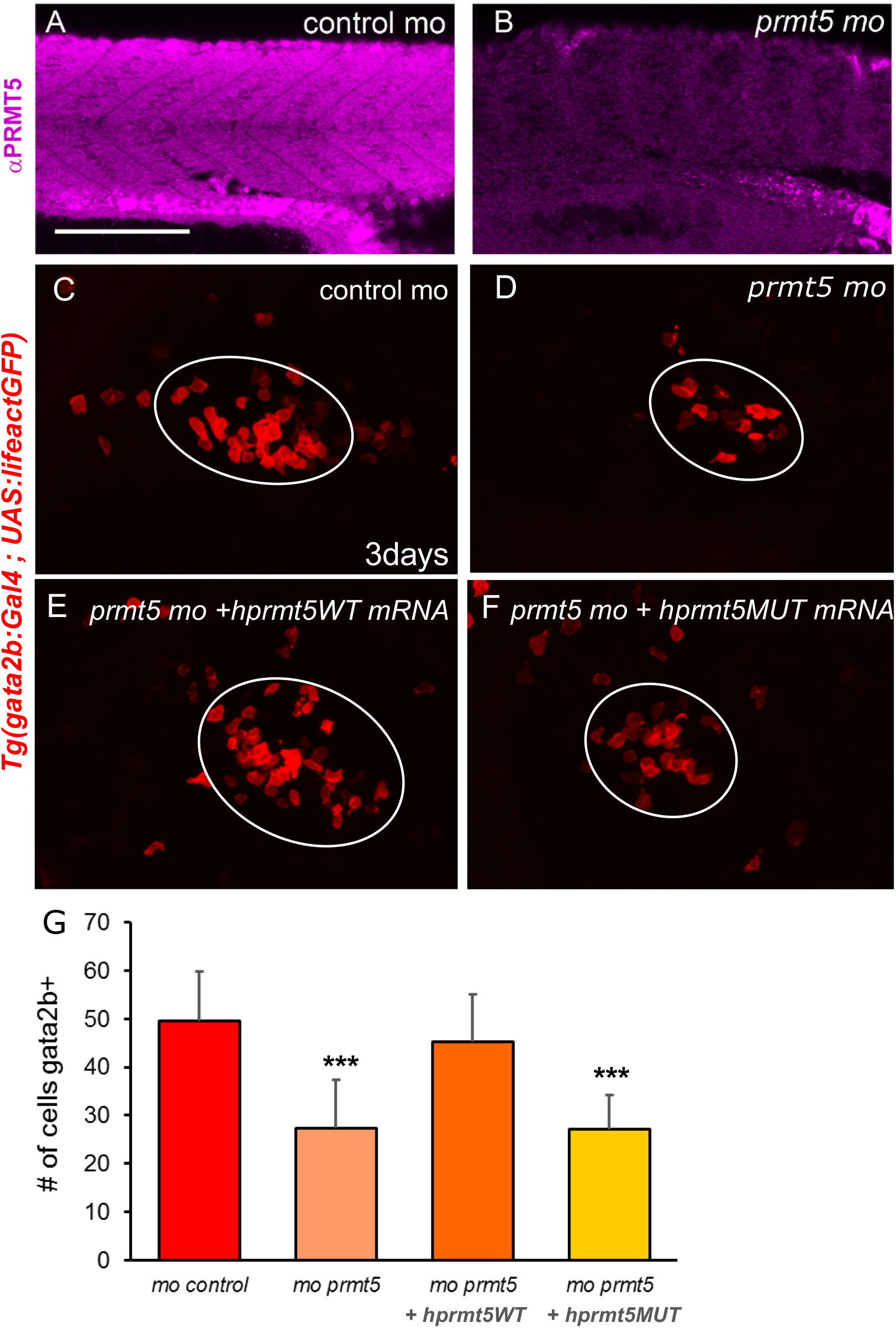
**A**-**B**-Confocal sections of Prmt5 immunostaining in control and *prmt5* morphant embryos at 24 hpf. Scale bar 100 μm. **C**-**F**-Confocal sections of thymus from transgenic Tg(*gata2b:Gal4; UAS:lifeactGFP*) embryos at 3 days. Transgenic embryos were injected by control morpholino (**C**) or *prmt5* morpholino only (**D**) or in combination *hprmt5WT* mRNA (**E**) or the catalytic mutant form *hprmt5MUT* mRNA (**F**). Thymus is delimited by a white circle. Bar scale 100 μm. **G**-Average number of HSPCs enumerated per confocal stack in injected embryos at 3 days from 2 independent experiments with at least 3 individuals per analysis. T-test was performed. ***P<0.001.

**Figure S2:**
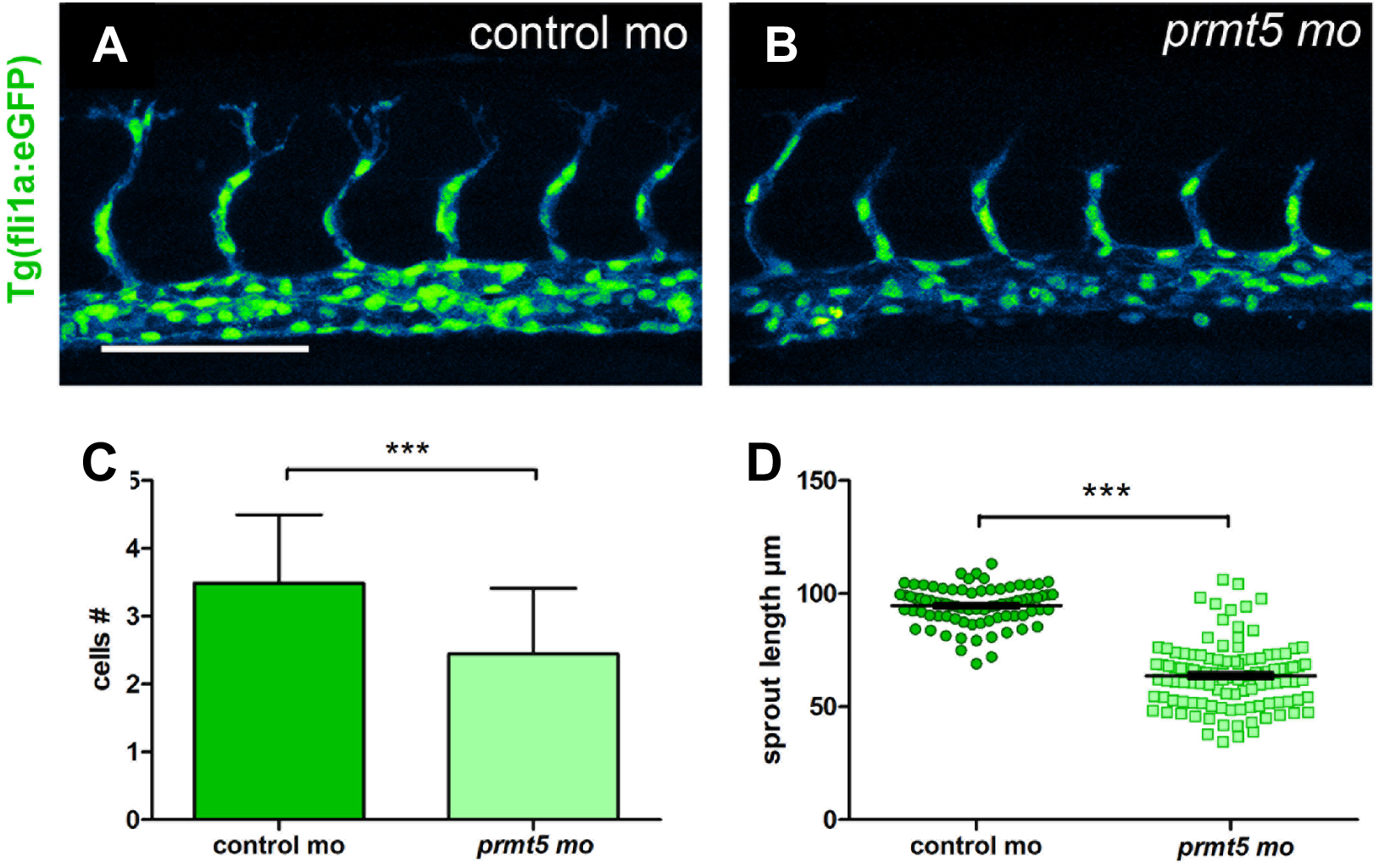
**A**-**B**-Confocal projections of transgenic Tg(*fli1a:GFP*)^y1^ embryos injected by either control morpholino (**A**) or *prmt5* morpholino (**B**). Scale bar 100 μm. **C**-**D**-Average number of endothelial cells per ISV (**C**) and average ISV length in μm (**D**), in control and *prmt5* morphant embryos, from 3 independent experiments with at least 4 animals per condition. T-test and Mann-Whitney test were performed. *** P<0.001.

**Figure S3:**
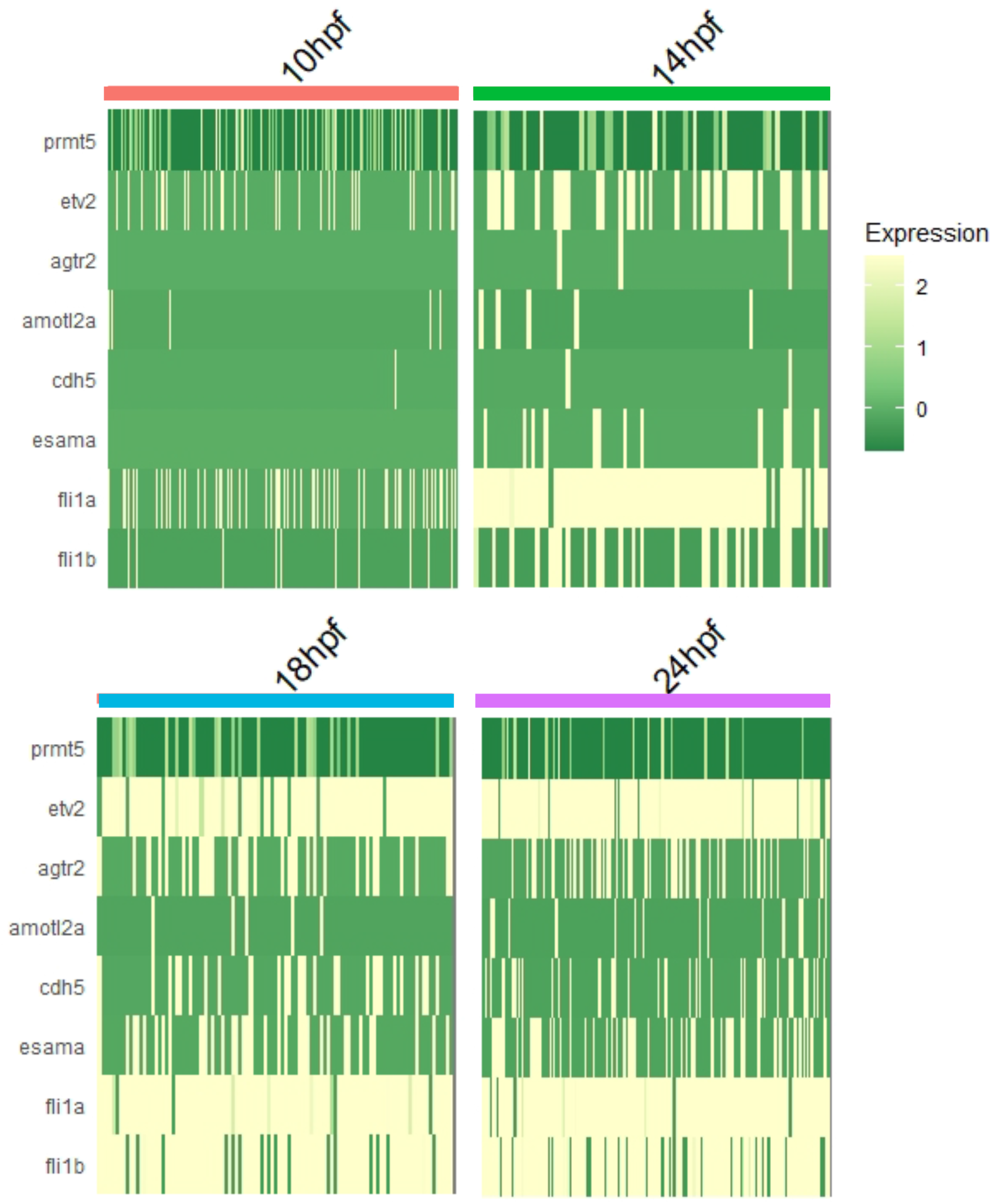
Expression heatmap for *prmt5*, *etv2* and identified Prmt5 target genes, for endothelial cells at 10hpf, 14hpf, 18hpf and 24hpf. The expression level is colored-coded from absence of expression (in green) to highest level of expression (in white).

**Figure S4:**
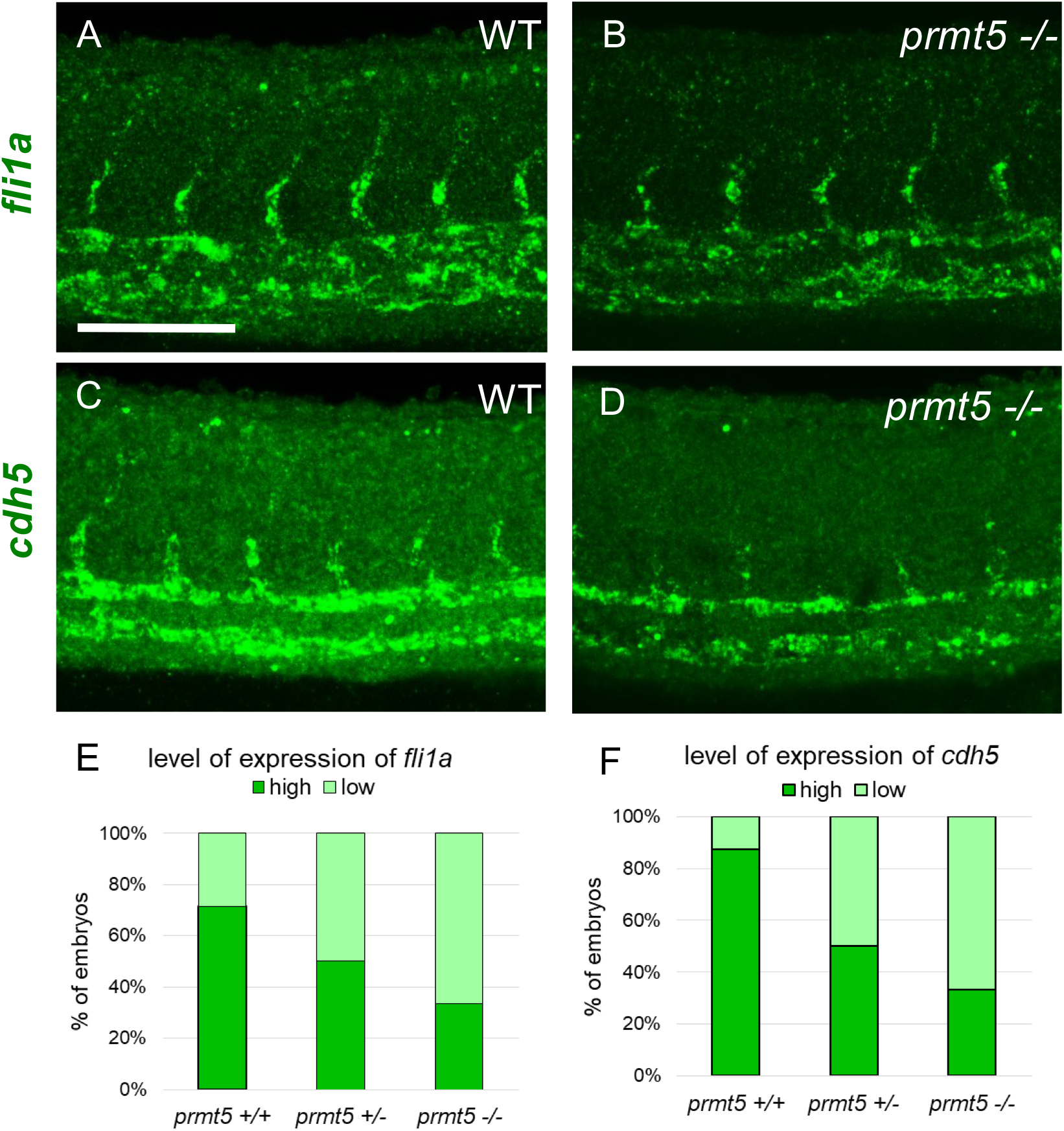
**A**-**D**-Confocal projections of wild type (**A**, **C**) and *prmt5* mutant embryos (**B**, **D**) after fluorescent *in situ* hybridization against *fli1a* (**A**-**B**) and *cdh5* (**C**-**D**). Scale bar 100 μm. **E**-**F-** Percentage of embryos (y axis) presenting a high or a low level of expression of *fli1a* (**E**) or *cdh5* (**F**), according to their genotype (x axis), from 3 independent experiments with at least 4 animals per condition.

**Figure S5:**
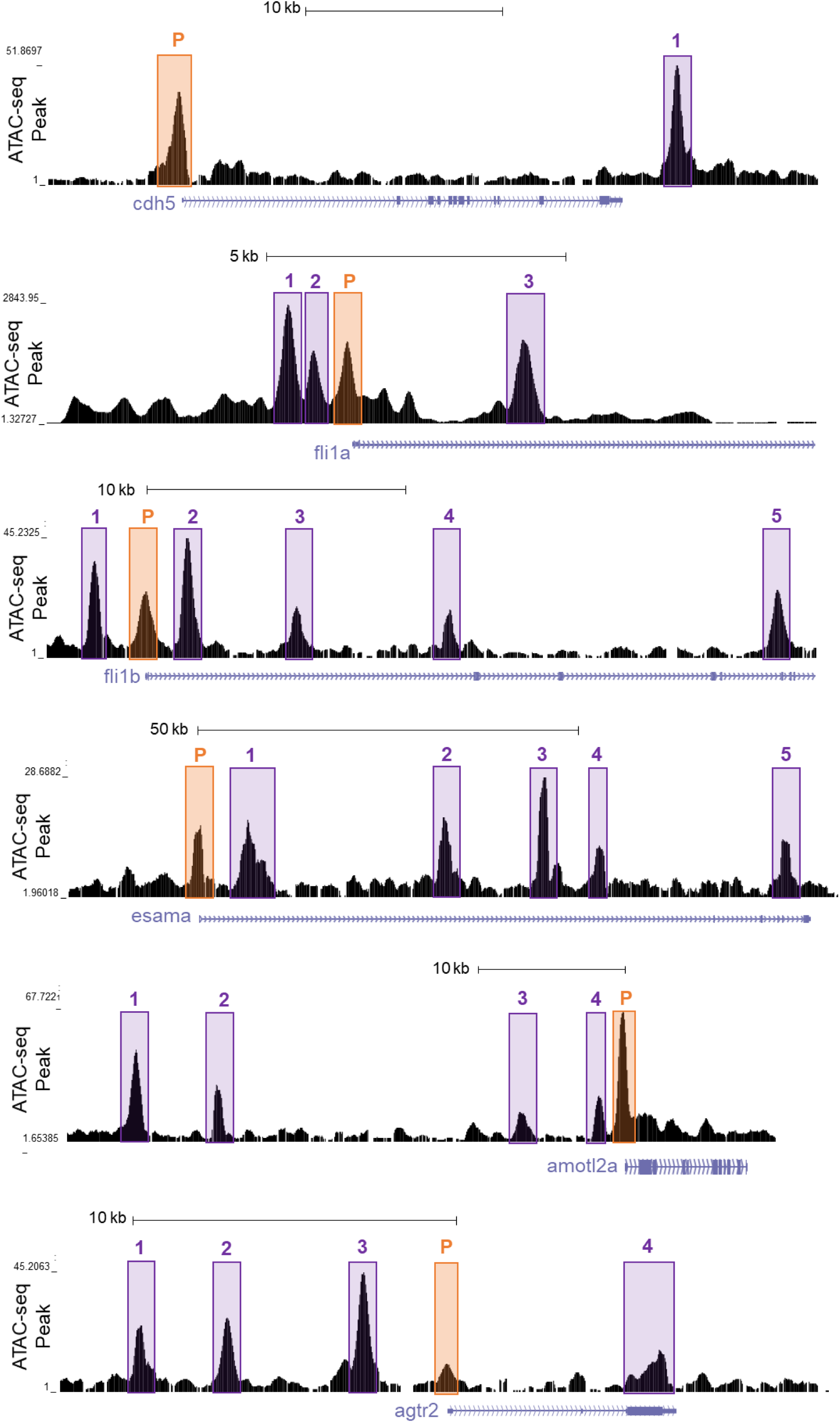
Chromatin profile visualization of endothelial cells from the UCSC Genome Browser. ATAC-seq peaks as determined by Quillien et al. (Quillien et al., 2017) flanking indicated genes (*cdh5, esama, agtr2, fli1a, fli1b, amotl2a*). Promoter regions (P) and numerated putative enhancers (corresponding numerated peaks are found in Table S1) are highlighted in light orange and light purple, respectively.

**Table S1.** Chromatin profile of Prmt5 identified target genes in zebrafish. ATAC-seq identified open chromatin regions surrounding characterized Prmt5 target genes and their distance to corresponding TSS (Quillien et al. 2017).

| Gene symbol | Peak chr | Peak start | Peak stop | ATAC-seq peak ID (fig. S5) | Distance of peakcenter to TSS |
| --- | --- | --- | --- | --- | --- |
| *fli1a* | *chr18* | 46991096 | 46991446 | 1 | 1 125 |
| *fli1a* | *chr18* | 46991648 | 46991909 | 2 | 663 |
| *fli1a* | *chr18* | 46992111 | 46992384 | *p* | 0 |
| *fli1a* | *chr18* | 46994744 | 46995457 | 3 | 2 866 |
| *fli1b* | *chr16* | 44785440 | 44785639 | 1 | 1 971 |
| *fli1b* | *chr16* | 44787371 | 44787644 | *p* | 0 |
| *fli1b* | *chr16* | 44788987 | 44789387 | 2 | 1 586 |
| *fli1b* | *chr16* | 44793201 | 44793495 | 3 | 5 851 |
| *fli1b* | *chr16* | 44799123 | 44799348 | 4 | **11 640** |
| *fli1b* | *chr16* | 44811618 | 44812080 | 5 | **24 462** |
| *esama* | *chr10* | 32600339 | 32600842 | *p* | 0 |
| *esama* | *chr10* | 32607536 | 32607916 | 1 | 6 894 |
| *esama* | *chr10* | 32632710 | 32633102 | 2 | **32 357** |
| *esama* | *chr10* | 32645662 | 32646247 | 3 | **45 230** |
| *esama* | *chr10* | 32653326 | 32653687 | 4 | **52 511** |
| *esama* | *chr10* | 32677711 | 32677978 | 5 | **76 617** |
| *amotl2a* | *chr6* | 27670527 | 27671187 | 1 | **29 519** |
| *amotl2a* | *chr6* | 27675738 | 27676044 | 2 | **24 324** |
| *amotl2a* | *chr6* | 27693946 | 27694363 | 3 | 6 172 |
| *amotl2a* | *chr6* | 27698667 | 27698822 | 4 | 1 629 |
| *amotl2a* | *chr6* | 27699920 | 27700375 | *p* | 0 |
| *cdh5* | *chr7* | 45432986 | 45433605 | *p* | 0 |
| *cdh5* | *chr7* | 45458271 | 45458821 | 1 | **24 800** |
| *agtr2* | *chr5* | 24901724 | 24902030 | 1 | **9 766** |
| *agtr2* | *chr5* | 24904437 | 24904720 | 2 | 6 990 |
| *agtr2* | *chr5* | 24908159 | 24908311 | 3 | 3 233 |
| *agtr2* | *chr5* | 24908459 | 24909026 | 4 | 2 645 |
| *agtr2* | *chr5* | 24917551 | 24917707 | *p* | 0 |
| *agtr2* | *chr5* | 24917756 | 24918031 | 5 | 6 525 |

**Table S2.**
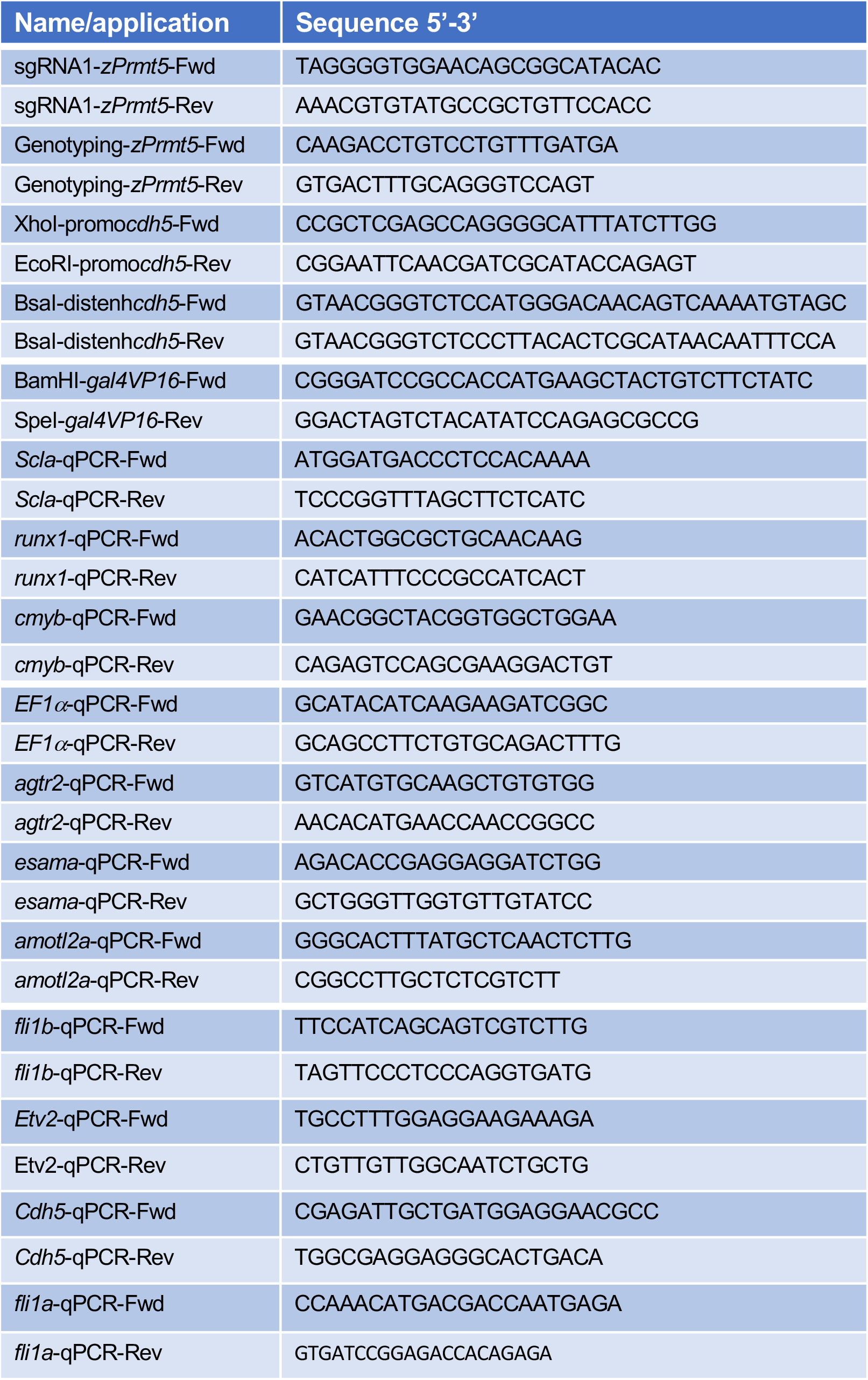
Sequences of oligonucleotides used in this study.

## Notes

### Competing Interest Statement

The authors have declared no competing interest.

